# Big Defensin *Ap*BD1 from the scallop *Argopecten purpuratus* is an antimicrobial peptide which entraps bacteria through nanonets formation

**DOI:** 10.1101/2020.02.25.965327

**Authors:** Felipe Stambuk, Claudia Ojeda, Paulina Schmitt

## Abstract

Antimicrobial peptides (AMPs) are ancient innate immune components. Big defensins is a family of AMPs found in a restricted number of animal phyla, in particular mollusks where they have highly diversified. Big defensins are composed of a highly hydrophobic N-terminal region and a C-terminal β-defensin-like region, stabilized by three disulfide bridges. They have been shown to be active against both Gram-positive, Gram-negative bacteria and fungi. Antimicrobial aggregates called nanonets entrapping bacteria have been recently described as the mechanism of action of the *Cg*-BigDef1 from the oyster *Crassostrea gigas*. Specifically, the N-terminal domain of *Cg*-BigDef1 was identified as responsible of nanonet formation. In order to determine whether nanonets are specific to oyster *Cg*-BigDef1 or common to other big defensins outside this species, we assessed the potential entrapping of bacteria through nanonets of the big defensin from the scallop *Argopecten purpuratus*, namely *Ap*BD1. Recombinant *Ap*BD1 was produced as a fusion polypeptide which carried a N-terminal His6 tag, with a thrombin cleavage site before the mature peptide sequence and an unfolded C-terminal domain by mutating the last Cys to Arg. Activity of r*Ap*BD1 was assessed against the gram-positive bacteria *Staphylococcus aureus* SG511. r*Ap*BD1 inhibited bacterial growth. Moreover, strong immune staining of r*Ap*BD1 in numerous areas surrounding bacteria was observed. Overall, results suggest that r*Ap*BD1 entrap bacteria in peptide aggregates similar to those reported to *Cg*-BigDef1. This study demonstrates the conservation of nanonet formation across big defensins and supports further a role for the N-terminal domain in this conserved process.

## 1. Introduction

Big defensins are a family of cationic antimicrobial peptides (AMPs) which are essential components of innate immunity. Big defensins are found in a restricted number of phyla, such as lophotrochozoans, certain arthropods and basal chordates (Loth *et al.* 2019). Big defensins are composed of a highly hydrophobic N-terminal region and a cationic C-terminal region containing six cysteine residues involved in three internal disulfide bridges (Rosa *et al.* 2011). The C-terminal domain of big defensins resembles β-defensins; recently they have been proposed to be the ancestors of this AMP family widespread in vertebrates (Zhu and Gao 2013). In certain species of mollusks, such as oysters and mussels, big defensins show high sequence diversity (Rosa *et al.* 2011; Gerdol *et al.* 2012). In contrast, only one isoform of the big defensin family has been identified so far in scallops (Zhao *et al.* 2007; Yang *et al.* 2016; González *et al.* 2017).

Studies related to the antimicrobial activities of big defensins have included assays testing the native polypeptide isolated from the horseshoe crab hemocytes (Saito *et al.* 1995), recombinant polypeptide produced in bacteria (Teng, Gao and Zhang 2012) and yeast (Zhao *et al.* 2007), and synthetic peptides obtained by chemical native ligation (Loth *et al.* 2019). In those studies, all big defensins showed antibacterial activity against Gram-positive, Gram-negative bacteria and fungi. The mechanism of action of the *Cg*-BigDef1 from the oyster *Crassostrea gigas* was recently identified. *Cg*-BigDef1 was found to entrap and kill *Staphylococcus aureus* in antimicrobial nanonets without inducing membrane permeabilization (Loth *et al.* 2019). Remarkably, the N-terminal domain of *Cg*-BigDef1 was the responsible of those fibrillar aggregation forming nanonets. Due to difficulties in producing sufficient amounts of pure big defensins, the horseshoe crab *Tt*-BigDef (Saito *et al.* 1995) and the oyster *Cg*-BigDef1 are the only members of the family for which significant data have been obtained in terms of mechanism of action. The recent discovery of bacterially-triggered nanonet formation in *Cg*-BigDef1 (Loth *et al.* 2019) raises the question of the conservation of this mechanism across big defensins.

With this objective we produced *Ap*BD1, the only big defensin found in the scallop *Argopecten purpuratus* previously studied in our research group (González *et al.* 2017). A polyclonal antibody was produced in mice against the mature peptide domain of the recombinant *Ap*BD1, which carried a N-terminal His6 tag followed by a thrombin cleavage site in frame with the mature peptide sequence, containing a mutation at the cationic C-terminal region. In the mutated polypeptide, the last Cys residue was replaced with an Arg residue by directed mutagenesis. As the recombinant *Ap*BD1 was produced to serve as antigen for antibody production, the disruption of the Cys stabilized structure was thought to improve the obtention of specific antibodies as previously suggested (Tykhomyrov *et al.* 2017).

Considering that the N-terminal domain of recombinant *Ap*BD1 (r*Ap*BD1) was integral and the nanonet formation by *Cg*-BigDef1 was related to the N-terminal domain of this AMP, we aimed to assess the potential nanonet formation by r*Ap*BD1. To do this, we cleaved the N-terminal His6 tag from the produced r*Ap*BD1 by thrombin cleavage and purified r*Ap*BD1 until homogeneity by RP-HPLC. When pure r*Ap*BD1 was incubated with the Gram-positive *S. aureus*, bacterial growth was disturbed by the polypeptide and similar images of strong immune staining of aggregates surrounding bacteria were found by immunofluorescence and confocal analysis as described for *Cg*-BigDef1. Results suggest that *Ap*BD1 from scallop also form nanonets to entrap bacteria as oyster big defensin.

## 2. Material and Methods

### 2.1 Recombinant expression and purification of rApBD1

The coding sequence for the mature peptide of *Ap*BD1 (GenBank No ANK58567) carrying a Cys to Arg substitution at position 121 of the complete amino acid sequence was cloned in-frame with the N-terminal His6 tag in the NheI/NdeI sites, which precedes a thrombin cleavage site on the pET-28a expression vector (Novagen). The recombinant His6-r*Ap*BD1 was expressed in *E. coli Rosetta* (DE3) and produced as previously described (González *et al.* 2017). Recombinant *Ap*BD1 was expressed at 37 °C in Luria Bertani broth by induction with 0.1 mM isopropyl-β-D-1-thiogalactopyranoside for 4 h at 37 °C and bacteria were lysed by sonication in 6 M guanidine-HCl in 100 mM Tris-Cl (pH 8.1). The His-tagged fusion peptide was purified using nickel–nitrilotriacetic acid (Ni–NTA, Qiagen) resin affinity chromatography and by C18 reverse-phase chromatography (Sep-Pak^®^ Waters) using a step gradient of 5% and 80% of acetonitrile; ACN, in acidified water with 0.05% trifluoroacetic acid; TFA). Fractions were concentrated by rotoevaporation in a SpeedVac Concentrator Savant SPD1010 (Thermo Scientific) and lyophilized. The elimination of the N-terminal His tag was performed through the cleavage with biotinylated thrombin included in the Thrombin Cleavage Capture Kit (Merck, Millipore) following manufacturer’s instructions. Thrombin was removed by incubating the solution with streptavidin agarose beads included in the kit. Cleaved peptide was purified by RP-HPLC (Jasco Analytical Instruments) with an Atlantis^®^ dC18, 3 μm × 4.6 mm × 150 mm (Waters) column. using a gradient of 5% to 60% ACN in 0.05% TFA in 60 min at a flow rate of 1 ml/min. UV absorption was read at 214 nm. HPLC fractions were concentrated and lyophilized. The molecular mass of purified peptide was determined by 18% SDS-PAGE and the peptide concentration was obtained by Bicinchoninic protein assay kit (Thermo Scientific).

### 2.2 Bacteria-rApBD1 interaction assay and Immunofluorescence

*S. aureus* SG511 was cultured overnight in Luria Bertani broth at 30 °C. Bacteria were washed three times by centrifugation and addition of killing buffer (KB; 0.1 M Tris [pH 8.0], 1 mM CaCl2). Bacterial concentration was adjusted to 10^7^ CFU/ml and 90 μl of bacterial suspension was incubated with 10 μl of purified r*Ap*BD1 (5 μM final concentration). Bacterial suspension incubated with 10 μl of KB, and buffer with peptide alone were also included as controls. Four replicates were performed for each condition. Two replicates of each condition were incubated over clean and sterilized coverslips in a 24-well plate for 18 h at 25 °C. The other two replicates were incubated in Eppendorf tubes at 25°C and after 4 h bacterial suspensions were centrifuged onto glass slides (10 min, 1,500 rpm, room temperature). All replicas were fixed for 10 min in 4% paraformaldehyde in PBS. Immunofluorescence analysis was carried out as described previously using the polyclonal antibody specific for ApBD1 (González *et al.* 2017). Bacteria were permeabilized with 0.2% Triton X-100 in PBS for 20 min and blocked for 2 h at room temperature with 3% PBSA (Bovine Serum Albumin in PBS). Then, slides were incubated overnight at 4°C with anti-ApBD1 (1:100) in 1% PBSA. A second incubation was performed for 1 h at room temperature with anti-mouse Alexa Fluor 568-conjugated (Thermo Scientific) 1:200 in 1% PBSA and then incubated for 5 min with Syto9 (1:1000) ^®^ (Thermo Scientific) for nuclear staining. Control slides were incubated with the prebleed serum of mouse. The slides were then mounted in Fluoromount mounting media (Sigma) and analyzed using a Leica TCS SP5 II spectral confocal microscope (Leica Microsystems). The images were obtained with a Leica 40×1.25 Oil HCX PL APO CS lens (Leica Microsystems).

## 3. Results and Discussion

### 3.1 rApBD1 was obtained after thrombin cleavage and RP-HPLC purification

The produced His6-r*Ap*BD1 fusion protein display a molecular weight of 11540.9 Da, which included a N-terminal His6 tag joined to a thrombin cleavage site (Leu-Val-Pro-Arg-ll-Gly-Ser; where ll denotes the cleavage site), with a molecular mass of 1882 Da. **(Figure 1A).** After replacing the last Cys residue to an Arg residue, the polypeptide with a theoretical molecular weight of 11540.9 Da was strongly induced in *E. coli*, which allowed us to partially purify His6-r*Ap*BD1 by affinity chromatography **(Figure 1B, lane 1)**. After obtaining the partial pure fraction, the cleavage of the His6 by thrombin allowed the recovery of a polypeptide with a molecular weight around 10000 Da **(Figure 1B, lane 2)**. The theoretical molecular weight of rApBD1 is 9658 Da, which is very close to the molecular weight of the bands observed by the SDS PAGE. After the removal of the biotinylated thrombin with streptavidin agarose beads, the resolution of the band around 10000 Da contained in the analyzed fraction was improved **(Figure 1B, lane 3)** and therefore, suitable for RP-HPLC further purification.

**Figure 1.**
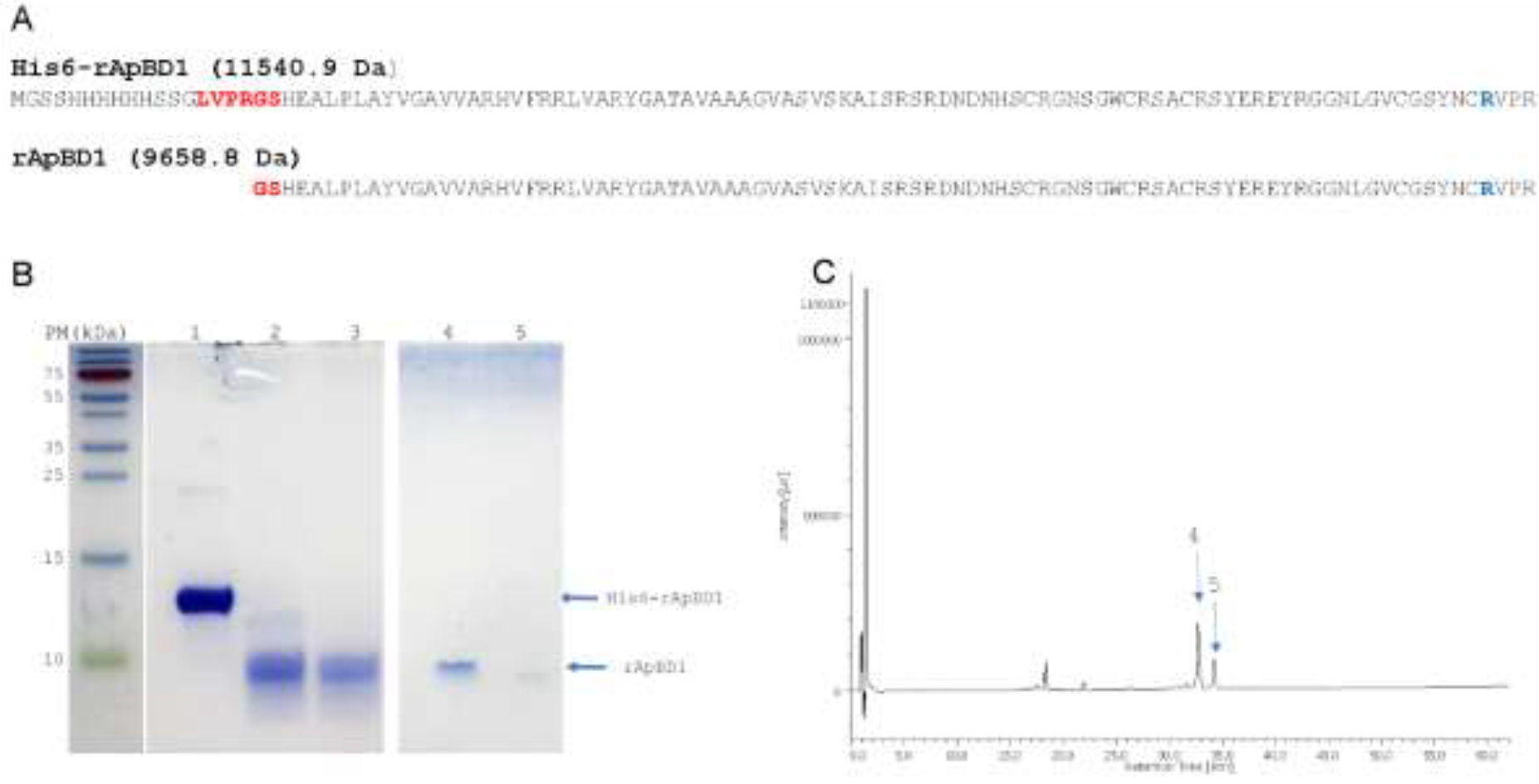
**A.** Amino acid sequences of fusion recombinant polypeptide His6-rApBD1 and cleaved rApBD1. Theoretical molecular weight is indicated (Da) for each sequence. Amino acids in red indicates thrombin cleavage site. In blue, Cys to Arg mutation introduced in the original sequence. **B.** SDS-PAGE showing purification steps of rApBD1. Lane 1. His6-rApBD1 after affinity chromatography. Lane 2: rApBD1 after thrombin cleavage. Lane 3. rApBD1 after thrombin removal. Lane 4 and 5. Two main fractions obtained by RP-HPLC. **C.** RP-HPLC profile of cleaved rApBD1.Fraction 4 and 5 are resolved in panel B.

After the RP-HPLC purification step, two main fractions were eluted around 30% ACN (between 32 and 35 min), namely fractions 4 and 5 **(Figure 1C)**. Those fractions were analyzed by SDS PAGE, showing the enrichment and purity of fraction 4 as r*Ap*BD1 **(Figure 1B, lane 4 and 5)**. The single band resolved in the SDS PAGE means a highly pure polypeptide and therefore we then assessed the potential interaction of ApBD1 with the bacteria *S. aureus* SG511.

### 3.2 *rApBD1* aggregates, entraps and inhibits the growth of *S. aureus*

Next, we deepen the interaction between r*Ap*BD1 and bacteria. As the β-defensin like C-terminal domain was not properly folded in r*Ap*BD1, we did not perform the standard antimicrobial assays used to determine the minimal inhibitory concentrations of cysteine-stabilized AMPs (Yang *et al.* 2000; Seo *et al.* 2005; Gueguen *et al.* 2009). However, r*Ap*BD1 possesses an integral N-terminal domain enabling to study the potential aggregation properties of this peptide and compare them to the bacterially-triggered formation of nanonets that was described for *Cg*-BigDef1 (Loth *et al.* 2019). We therefore assessed the potential formation of r*Ap*BD1 aggregates upon contact with bacteria. When rApBD1 (5 μM) was added to a *S. aureus* monolayer culture for 18h, we observed that the growth the bacterial cells was altered, with lower numbers of adherent bacteria and more visible bacterial clumps **(Figure 2A, right panel)**. In addition to growth inhibition, we observed a strong immune staining of r*Ap*BD1 in numerous extents surrounding bacteria **(Figure 2B),** as earlier described for *Cg*-BigDef1 nanonets in contact with *S. aureus* (Loth et al, 2019). No immune staining was detected in control samples in which no r*Ap*BD1 was added to bacteria, and no aggregation of the peptide were observed in the absence of *S. aureus* (data not shown). When confocal sections were analyzed, images revealed no immune staining at bacterial cells place, suggesting that r*Ap*BD1 did not lyse bacteria but rather entrapped them in aggregates similar to those reported to CgBigDef1 (Loth *et al.* 2019). In *C. gigas*, the N-terminal domain alone also produced nanonets but did not kill *S. aureus*. This led the authors to hypothesize that *Cg*-BigDef1 antimicrobial activity is carried by the β-defensin-like domain, but needs the N-terminal domain to get in contact with microorganisms (Loth *et al.* 2019). Our present data support further this hypothesis since the C-terminal β-defensin-like domain of the r*Ap*BD1 produced in this study lacks a Cys residue. It is therefore very likely that the intact N-terminal domain of the molecule is responsible for the nanonet formation.

**Figure 2.**
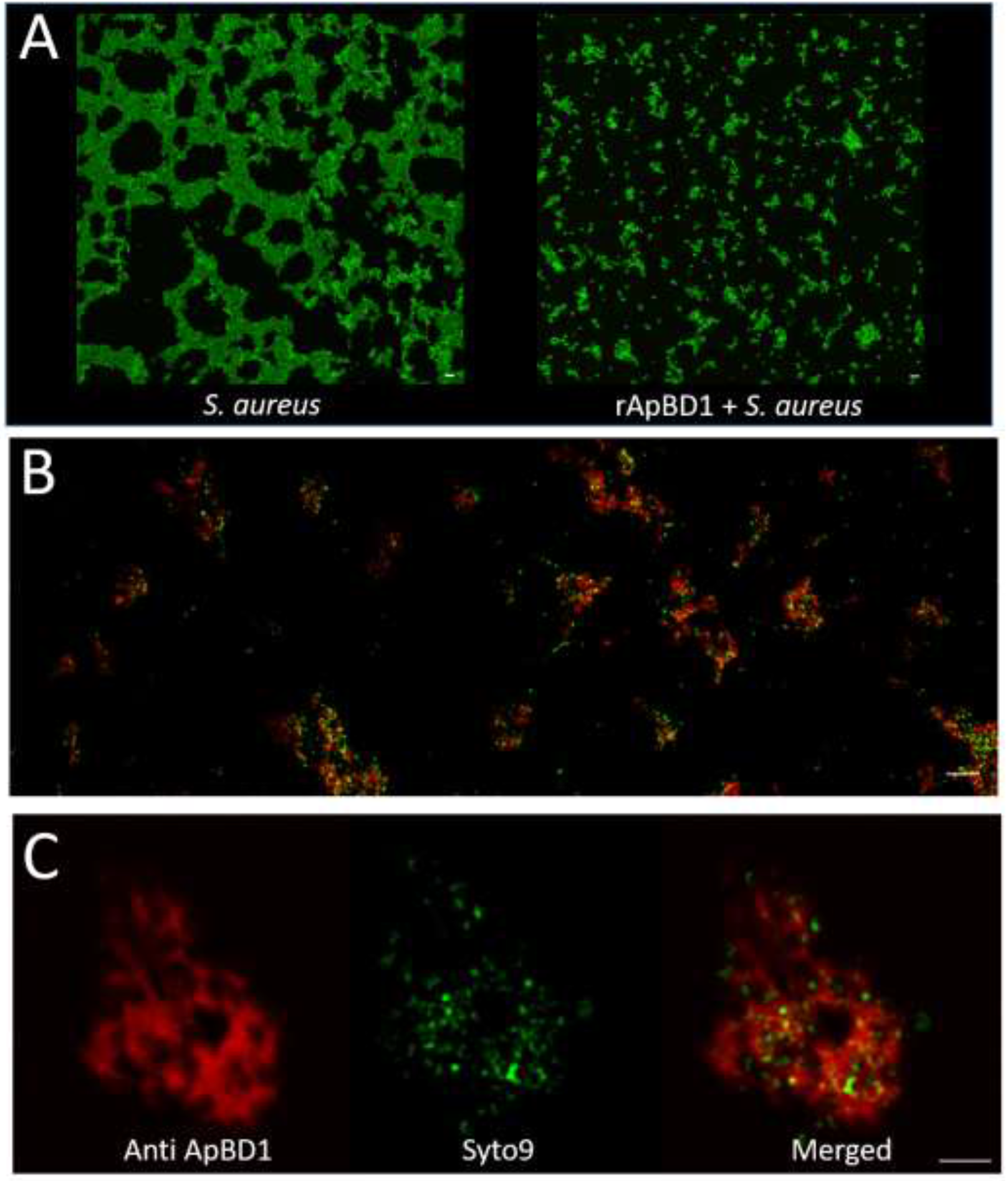
Interaction between rApBD1 and *S. aureus* by immunofluorescence and confocal analysis. **A.** Monolayer culture of *S. aureus* for 18 h at 25°C. Left panel, *S. aureus* alone. Right panel, *S. aureus* with 5 μM of rApBD1. Bacterial DNA was stained with Syto9^®^ (green). **B.** General view of immune staining of ApBD1 incubated with *S. aureus* for 4 h at 25°C. Bacterial DNA was stained with Syto9^®^. rApBD1 was stained with anti-ApBD1 and anti-mouse Alexa Fluor 568-conjugated (red). **C.** Confocal analysis of observed aggregates. Merged image showed *S. aureus* cells entrapped in rApBD1 aggregates similar to nanonets. Scale bars: 3 μm.

Altogether our data show that nanonet formation is not restricted to big defensins from *C. gigas* oysters but is conserved in rather distant species of bivalve mollusks like the scallop *A. purpuratus*. In line with the formal demonstration made in oyster (Loth et al., 2019), the N-terminal domain conserved in big defensins could play a key role in *Ap*BD1 mechanism of action.

### Concluding Remarks

Scallop rApBD1 is a member of the big defensin family which are capable of entrap and impair normal growth of *S. aureus* as previously observed for the Cg-BigDef1 from the oyster. In the future, the native sequence must be obtained assess further mechanisms of action of this important scallop AMP.

